# Activity-level evaluation of DNA transfer to stockings: comparison of HaloGen with simplified contributor-based approaches

**DOI:** 10.64898/2026.09.04.749444

**Authors:** Peter Gill, Helen Johannessen, Christine Børsum, Ane Elida Fonneløp, Øyvind Bleka

## Abstract

DNA findings on clothing are often evaluated at sub-source level to address whether a person contributed DNA to a sample. In many cases, however, the disputed issue is not whose DNA is present, but how the DNA was transferred. This paper applies the HaloGen framework to evaluate DNA findings on stockings under activity-level propositions. The data derive from controlled experiments, reported in a companion paper, comparing active contact, in which a person of interest pulled down a pair of stockings, with social contact, in which the person of interest and wearer used the same bathroom. Contributor-specific DNA quantities were estimated for the person of interest and for observed unknown contributors across two sampled stains.

Compatibility diagnostics indicated that the stockings dataset was broadly compatible with the observed Pop20 reference range. Pop20 was therefore used as regularising prior information in the Lab-Bayes model, rather than as a stand-alone Group model.

Because the stockings dataset was generated in a single laboratory, transfer data from 20 laboratories were used as regularising prior information for a laboratory-specific Lab-Bayes model. The local stockings data then updated the direct- and secondary-transfer parameter distributions. Compatibility diagnostics indicated that the stockings dataset was broadly compatible with the observed Pop20 reference range. Pop20 was therefore used as regularising prior information in the Lab-Bayes model, rather than as a stand-alone Group model. In addition, prior-sensitivity and leave-one-case-out analyses indicated that the case-level likelihood ratios were not materially affected by the choice of prior specification or by data reuse.

HaloGen likelihood ratios were assessed using Tippett plots and compared with simplified contributor-based comparator approaches. These approaches represent simplified forms of activity-level reasoning in which the DNA result is reduced to binary or categorical states. They comprise a binary source-support approach, a discrete *M_x_* component-size approach, and a background approach based on the ReAct I Experiment 3 formulation. These approaches were used to examine the effect of reducing continuous contributor-specific DNA quantities to binary or categorical states, and to assess how the likelihood ratio is affected when observed unknown contributors are represented through a background/direct-unknown term rather than evaluated explicitly as possible alternative direct actors.

HaloGen showed clear discrimination between active-contact and social-contact ground-truth cases. The simplified comparator methods illustrated the information loss caused by thresholding or categorising continuous DNA quantities. The ReAct background approach tended to shift direct-transfer likelihood ratios upward relative to the HaloGen calculation, showing that the treatment of observed unknown contributors can materially affect activity-level inference. The results support the importance of preserving contributor-level symmetry: an observed unknown contributor should not automatically be demoted to background when that contributor may represent an alternative direct actor under the defence proposition.

## 1 Introduction

This investigation was partly motivated by the Birgitte Tengs case [9]. DNA findings on clothing are frequently encountered in cases where the source of the DNA is not disputed, but the mechanism of transfer is. In such cases, the relevant issue is not simply whether DNA from a relevant actor, such as the POI, is present, but whether the observed DNA quantities are more probable under one activity proposition than another. For example, the prosecution may allege that the POI directly handled or removed a garment, while the defence may propose that the DNA was transferred indirectly or innocently through social contact [14, 25, 2].

The waistband and hip region of stockings, trousers, or underwear is a common target area in alleged sexual assault investigations because it may correspond to the area grasped during removal of the garment. However, DNA recovered from such areas may include the wearer, the POI, and one or more unknown contributors. The presence of unknown contributors raises a central interpretive question: should those contributors be treated as background, or should they be evaluated as possible alternative direct actors under the defence proposition?

This distinction is important at activity level. If an observed unknown contributor is automatically reduced to background, the model may effectively privilege the named POI as the only contributor eligible to occupy the direct-transfer role. The resulting likelihood ratio may then favour the prosecution proposition for a structural reason rather than an evidential one because the model structure has prevented the unknown contributor from competing as an alternative direct actor. A symmetry-preserving model should instead allow observed unknown contributors to be evaluated explicitly. This is consistent with broader work on exhaustive treatment of multiple persons of interest in DNA mixtures and with case-based concerns about confirmation bias when unknown DNA is not properly considered [17, 23, 9].

HaloGen is a Bayesian activity-level framework developed to evaluate DNA quantities under competing transfer propositions [11, 10]. It uses experimental data to estimate direct-transfer and secondary-transfer distributions, and then evaluates contributor-specific quantities within an exhaustive assignment framework. The framework allows multiple contributors, multiple stains, observed unknown contributors, and the possibility that a relevant actor leaves no detectable DNA, represented by the non-detection probability *F*_0_.

The present paper applies HaloGen to a controlled stockings transfer dataset [18]. The dataset compares active contact, where the POI pulled down a pair of stockings, with social contact, where the POI and wearer used the same bathroom but the POI did not directly handle the stockings. The competing propositions were therefore that the POI directly pulled down the stockings, or that the POI had only social contact with the wearer and did not directly handle the stockings.

The aim of this methodological paper is to compare the full quantitative HaloGen calculation with simplified comparator methods applied to the same contributor-specific data. Specifically, we compare HaloGen with a binary source-support approach, a discrete *M_x_* component-size approach, and a (ReAct I project [13] Experiment 3) background approach. These comparisons assess the effect of information loss when continuous contributor-specific quantities are simplified to binary or categorical observations, and how the LR changes when observed unknown contributors are treated as background rather than as possible alternative direct actors.

The experimental design, laboratory methods, sample-level sub-source results and unfiltered contributor-specific dataset are reported in the companion experimental paper [18]. The present paper focuses on the activity-level behaviour of HaloGen and on comparison with simplified contributor-based comparator methods. It also assesses whether this new stockings dataset can be analysed within a Bayesian hierarchical framework informed by the ReAct transfer data, by using the ReAct data as regularising prior information and verifying the effect of that choice through compatibility, prior-sensitivity and leave-one-case-out checks.

## 2 Data and activity-level propositions

### 2.1 Experimental dataset

The data derive from controlled experiments designed to compare DNA transfer to stockings during active contact and social contact [18]. In the active-contact experiment, the POI pulled down a pair of stockings. In the social-contact experiment, the POI and the wearer used the same bathroom, without the POI directly handling the stockings. Two DNA samples were collected from the waistband/hip region of each pair of stockings, corresponding to the sampled areas most relevant to the alleged activity.

For the activity-level analysis, contributor-specific DNA quantities were required for the POI and for any observed unknown contributors. The female wearer was expected to contribute DNA under both propositions and was therefore not evaluated as a competing relevant actor (i.e. simply omitted from analysis). The activity-level analysis focused on the POI and observed unknown contributors detected in the sampled stains.

For the current analysis, the paired activity-level dataset comprised 20 active-contact cases and 20 social-contact cases (40 cases total), each represented by two stain-level observations (80 stain-level observations total); see [18] for the experimental design and full sample-size accounting.

### 2.2 Contributor-specific quantity estimation

Contributor-specific quantities were obtained by combining total DNA quantity with contributor mixture proportions estimated at sub-source level, as described in the companion experimental paper [18].

Let *Q*_total_ denote the total DNA concentration in the extract, *E* the elution volume, and *m_i_* the mixture proportion assigned to contributor *i*. The estimated quantity for contributor *i* was

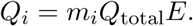

The elution volume was *E* = 200 *µ*L.

EFMrep [5, 4] was used because two STR profiles (an inside and an outside sample) were analysed for each activity. These two samples were modelled jointly under a common set of contributor mixture proportions, rather than estimating separate mixture proportions for each. This produced a combined sample-level mixture-proportion estimate for each contributor, which was then used to calculate the contributor-specific DNA quantity for the activity-level analysis. Sub-source propositions and replicate handling are described in the experimental paper [18]. Because the ground truth was known in the controlled experiments, the expected female donor and the POI were included in the sub-source modelling; where additional alleles could not be explained by these contributors, the number of contributors was increased to accommodate one or more unknown contributors.

Contributor-specific quantity construction and the full contributor-specific HaloGen likelihoodratio tables are provided in Supplementary Material S1.

### 2.3 Activity-level propositions and one-actor assumption

The activity-level propositions were:

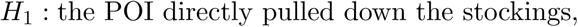

and

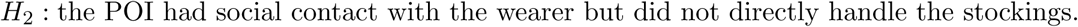

Here, a direct actor means the person assigned to the activity-relevant direct-contact role: the person who directly handled or pulled down the stockings. A secondary contributor is a person whose DNA may be present as a result of indirect transfer or social contact rather than direct handling of the stockings. In this example, under *H*_1_, the POI is the direct actor. Under *H*_2_, the DNA of the POI is assumed to originate from an indirect contact; an observed unknown contributor may instead be considered as the possible direct actor, or the relevant direct actor may be unobserved. This distinction is central to the HaloGen calculation because observed unknown contributors are not automatically treated as background.

## 3 HaloGen model and calibration

### 3.1 Overview of the HaloGen activity-level calculation

HaloGen models DNA quantities under direct and secondary transfer using zero-augmented and left-censored lognormal mixture models. For each transfer assumption, the distribution is described by parameters (*µ, σ, k*), where *µ* and *σ* define the lognormal component and *k* represents the structural non-transfer or dropout component. The detection limit was specified as *DL* = 0.001 ng, matching the lower censoring limit used for the quantity inputs.

For detected quantities, HaloGen uses the lognormal component conditional on detection. Thus, for *q > DL*, the case-level likelihood under a transfer model with lognormal density *f* (*q*) and cumulative distribution function *F* (*q*) is

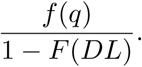

This is the density of a lognormal distribution left-truncated at the detection limit. Detected direct- and secondary-transfer quantities are therefore evaluated using the corresponding left-truncated lognormal densities.

Non-detection of a required direct actor is represented separately through *F*_0_, the probability that the relevant direct actor leaves no detectable DNA:

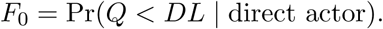

For each observed contributor, HaloGen evaluates the likelihood of the contributor’s quantity under direct transfer and secondary transfer. These likelihoods are then combined across stains and across admissible contributor-role assignments. Let *N_S_* denote the number of relevant direct actors; in this study *N_S_*= 1. In simplified notation, for an elemental assignment in which contributors in set *O* are direct actors and the remaining contributors are secondary, the elemental likelihood contains terms of the form

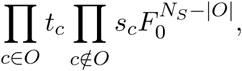

where *t_c_* and *s_c_* are the contributor-specific direct and secondary likelihoods, respectively. The *F*_0_ term accounts for any required direct actor who is not detected among the observed contributors. The final likelihood ratio is obtained by summing over all admissible elemental assignments within each top-level proposition, as described by [11]. Full formulae for the HaloGen likelihood construction are given in Supplementary Material S2.

### 3.2 Primary Lab-Bayes model informed by ReAct reference dataset

Hierarchical Bayesian models provide a natural way to partially pool information across related sources while retaining laboratory-specific estimates; this is particularly relevant where local calibration data are limited but related multi-laboratory data are available [1, 13, 8, 26]. In the present paper, the Pop20 dataset from the ReAct project [13] was used as external reference information for the direct- and secondary-transfer parameters. Pop20 denotes the 20-laboratory transfer dataset used to define the regularising prior information.

The stockings dataset was generated in a single laboratory. A stockings-specific multi-laboratory Group model could therefore not be estimated, because there was no collection of stockings experiments performed across multiple laboratories from which to estimate between-laboratory variation for this specific experimental system. The available Pop20 reference data were derived from ReAct transfer experiments, not from a multi-laboratory stockings experiment. A pure Pop20 Group result would therefore represent a generic multi-laboratory prediction rather than a laboratory-calibrated analysis of the stockings dataset. For this reason, Group-model likelihood ratios were not used as the primary activity-level results for the stockings analysis.

The primary analysis instead used the Pop20 dataset to inform the Lab-Bayes model based on the stockings data. In this model, the Pop20 reference data define regularising prior information for the direct- and secondary-transfer parameters. The local stockings observations were then used to update these Pop20-informed priors. The first diagnostic examined whether the target stockings dataset, was sufficiently compatible with the Pop20 reference structure for it to be used as regularising prior information. This provides a regularised single-laboratory analysis: it does not replace the local stockings data, but stabilises inference where the local data alone are limited, particularly for the secondary-transfer distribution. The Pop20-informed Lab-Bayes model was therefore used in the final HaloGen model for the target stocking dataset, which was used for the method comparisons.

The suitability of using Pop20 as a source of prior information was assessed explicitly using compatibility diagnostics, prior-sensitivity analysis and leave-one-case-out analysis. These checks are summarised in Section 3.3, with detailed methods, tables and diagnostic plots provided in Supplementary Materials S3 and S4.

### 3.3 Diagnostics for the Lab-Bayes model informed by the ReAct Pop20 reference dataset

Three different kinds of model diagnostics were considered to assess whether the ReAct Pop20-informed Lab-Bayes model was suitable as the reference model for the method comparisons: representativeness relative to the Pop20 reference data, sensitivity to prior specification, and leave-one-case-out assessment of data reuse.

#### 3.3.1 Representativeness relative to the ReAct Pop20 reference data

The first diagnostic examined whether the target stockings dataset, was sufficiently compatible with the Pop20 reference structure for it to be used as regularising prior information. In this analysis we provided different parameter representations for investigating the difference of lab-specific parameters: Mahalanobis distances, principal component analyses and posterior predictive checks.

Different parameter representations were used to assess whether the stockings data were unusual relative to the Pop20 reference laboratories. The PCA plots showed substantial overlap between the target laboratory and the Pop20 laboratories in the first two principal components. The robust Mahalanobis diagnostics gave the same practical interpretation when assessed using empirical upper-tail fractions: the stockings target laboratory was not an isolated outlier within the finite Pop20 reference set, although it was not centrally representative in some absolute parameter representations. The posterior predictive checks showed good overlap for direct transfer and more limited overlap for secondary transfer at smaller quantities. Overall, the compatibility diagnostics indicated that the stockings data were within the empirical Pop20 reference range, although not centrally representative in all parameter representations. Pop20 was therefore used cautiously as prior information and updated by the local stockings observations, rather than used as a substitute for local data. Full compatibility methods, empirical upper-tail fractions, diagnostic tables and PCA plots are provided in Supplementary Material S3.

#### 3.3.2 Sensitivity to prior specification

The second diagnostic assessed whether the case-level results depended materially on the Pop20-informed prior specification. This used the same likelihood model, data structure, *F*_0_ policy, defined cases and case-level LR calculations as the primary analysis, but replaced the Pop20-informed prior with weakly informative hyperparameters that allowed substantially greater prior variance.

The direct-transfer posterior summaries were largely insensitive to the prior choices, indicating that the direct-transfer component was mainly data-driven. The secondary-transfer summaries showed greater prior sensitivity, as expected for the more variable and sparse secondary-transfer component. However, this did not materially affect the case-level conclusions. All weak-prior medians fell within the corresponding Pop20-informed 10–90% posterior intervals. The median weak-prior minus Pop20-informed shift was +0.21 log_10_ units for active-contact cases and +0.20 log_10_ units for social-contact cases. Full prior hyperparameters, posterior parameter summaries and case-level sensitivity plots are provided in Supplementary Material S4.

#### 3.3.3 Leave-one-case-out check

The third diagnostic assessed the effect of reusing the local stockings data for both calibration and case evaluation [24, 7, 28]. This issue is particularly relevant for the Tippett plots in Section 5.2, because the stockings cases are used as ground-truth test cases for assessing discrimination between active-contact and social-contact propositions. In the full-data Pop20-informed Lab-Bayes analysis, the same local stockings observations contribute both to the laboratory-specific transfer posterior and to the LR calculated for the case being plotted.

The leave-one-case-out procedure addressed this potential data-reuse issue by omitting the activity-level case being evaluated from the local calibration data and refitting the model using the remaining local data. The held-out case was then evaluated using the fold-specific posterior. The omitted unit was the activity-level case, not an individual stain; therefore, both stain observations belonging to the held-out case were excluded from the relevant local calibration arm. The external Pop20 prior information was kept fixed because it was estimated from data external to the stockings experiment.

This procedure provides a closer analogue of evaluating a case that did not contribute to the local calibration data. However, it requires a separate Lab-Bayes refit for each held-out case and means that different cases are evaluated using slightly different local calibration datasets. For the main Tippett plots and simplified-comparator analyses, the full-data Pop20-informed Lab-Bayes model was therefore retained as a convenient common reference model, provided that this approach did not materially affect the LR conclusions.

The leave-one-case-out results were very close to the full-data results: the median log_10_(*LR*) changed from 1.52 to 1.49 for active-contact cases and from *−*2.98 to *−*2.96 for social-contact cases. No case changed direction of support after leave-one-case-out recalibration. Detailed leave-one-case-out methods and diagnostics are provided in Supplementary Material S4.

Because the stockings laboratory was not centrally representative of Pop20, use of a pure Pop20 Group model as the primary result was not supported. Pop20 was therefore used only as regularising prior information in the Lab-Bayes model, with the local stockings observations providing the laboratory-specific update. The resulting case-level conclusions were robust to both weak-prior refitting and leave-one-case-out recalibration.

## 4 Comparison with discrete approaches

### 4.1 Purpose of the simplified comparator approaches

The simplified comparator approaches were not intended as alternative reporting models. They were included as controlled simplifications of the same evidence to examine the consequences of thresholding or categorising contributor-level observations. Binary or categorical approaches are analogous to discrete-state activity-level models in which the DNA result is represented by a categorical node rather than by continuous quantities [25, 20, 22, 3].

The full HaloGen calculation uses continuous contributor-specific DNA quantities and explicitly evaluates observed unknown contributors as possible direct actors. To assess the effect of simplifying this calculation, we compared HaloGen with three simplified contributor-based approaches:

1. a binary source-support approach described in Section 4.2;
2. a discrete *M_x_* component-size approach, described in Section 4.3; and
3. a ReAct3 background approach, described in Section 4.4.

The first two approaches use simplified binary or categorical observations but retain the same exhaustive contributor-assignment logic as HaloGen: observed unknown contributors are allowed to compete with the POI for the direct-actor role. The third approach also uses a discrete *M_x_*-based categorical representation, but replaces explicit observed unknown-contributor assignment with a background/direct-unknown term. This distinction is central to the comparison in Section 5.3.

The binary source-support and discrete *M_x_* approaches were implemented using the same exhaustive contributor-assignment logic as HaloGen. Work on logical frameworks for DNA interpretation, exhaustive propositions, and multiple persons of interest emphasises that propositions should be specified independently of the resulting LR and should account for relevant alternative contributor assignments [16, 17, 6, 23]. Full derivations of the binary source-support, discrete *M_x_* approach, and ReAct3 background approach are provided in Supplementary Material S5.

### 4.2 Binary source-support approach

For the binary source-support approach, contributor-level observations were simplified to binary states:

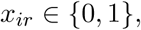

where *x_ir_*= 1 indicates that contributor *i* was treated as a supported contributor on stain *r*, and *x_ir_* = 0 indicates that the contributor was not treated as supported.

This was deliberately not a quantity-presence indicator based only on *Q_i_ > DL*. Instead, it represented a stringent sub-source support rule. Presence was defined using

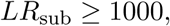

where *LR*_sub_ denotes the sub-source likelihood ratio obtained from the EFMrep analysis. The direct-transfer presence frequency was 35/40 and the secondary-transfer presence frequency was 0/40. Jeffreys correction was applied to avoid probabilities of zero or one, giving

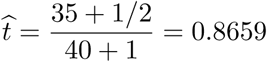

and

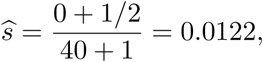

where *t* = Pr(presence *| D*) and *s* = Pr(presence *| S*).

The binary source-support approach should be interpreted only as a rough threshold-based benchmark. It discards continuous DNA quantity and mixture-proportion information, and therefore cannot reproduce the full stain-level information used by HaloGen.

### 4.3 Discrete ***M_x_*** component-size approach

The discrete *M_x_*approach classified each activity-relevant contributor on each stain as absent, minor, or major:

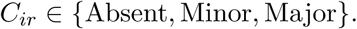

Major/minor terminology is well established in forensic DNA mixture interpretation, and propositions may be framed in terms of whether a named person or an unknown person is the major contributor [16]. A mixture-proportion threshold of approximately 0.6 has also been used previously as an operational definition of a major contributor in a related transfer and activity-level study [19]. For the present simplified comparator, the threshold was therefore defined operationally: a contributor was classified as absent if no DNA quantity was attributed to that contributor, as major if *M_x_ ≥* 0.6, and as minor if 0 *< M_x_ <* 0.6. The threshold was chosen to represent a clear majority component in this controlled comparison. It is deliberately restrictive and should not be interpreted as a general forensic standard; in mixtures with more contributors, a main component may have *M_x_ <* 0.6.

For the discrete *M_x_*approach, detected contributor assignments were retained only where

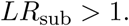

This was a minimum source-support restriction, not a reporting threshold. It retained assignments with at least some sub-source support while excluding assignments with no support for the contributor. Contributor-stain combinations with no attributed quantity were retained as absent observations for the purpose of estimating the absent category.

This restriction differs from the binary source-support threshold because the two approaches answer different simplified questions. The binary source-support approach asks whether a contributor would be treated as supported at a stringent sub-source level. The discrete *M_x_* approach asks whether a supported contributor is absent, minor, or major in component size. It retains more categorical information than the binary source-support approach, but it remains a simplified approximation and does not use the full contributor-specific quantities used by HaloGen.

### 4.4 ReAct Experiment 3 background approach

The third simplified approach used a variant of the ReAct Experiment 3 background approach described in [13], where the screwdriver was handled one hour after handshake. The purpose was not to reproduce the full ReAct analysis, but to examine the effect of replacing explicit unknown-contributor assignment by a background/direct-unknown term.

In this formulation, *H*_1_ considers the POI as the direct actor, while *H*_2_ considers the POI as secondary with an unknown direct actor. Let

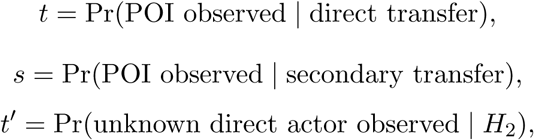

and

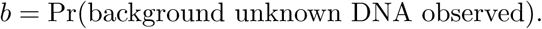

For a single stain where only the POI is observed, the ReAct3 expression is

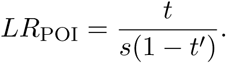

Where both the POI and an unknown contributor are observed, the corresponding expression is

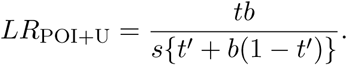

In the present analysis, the background parameter was set to *b* = 0.3.

The ReAct3 background approach is not equivalent to HaloGen because it does not evaluate each observed unknown contributor separately as a competing direct actor. It was therefore used as a simplified comparator to assess how a background-style treatment of observed unknown contributors affects the LR.

## 5 Results

### 5.1 Primary HaloGen activity-level likelihood ratios

The controlled stockings experiments were used for methodological comparison, with the full-data Pop20-informed Lab-Bayes analysis as the primary reference model.

The same dataset was used to compare output from simplified comparator approaches.

HaloGen generated contributor-specific activity-level likelihood ratios for the POI and for each observed unknown contributor. The female wearer was expected under both propositions and was not evaluated as a competing activity-relevant actor. The Pop20-informed Lab-Bayes median log_10_(*LR*) was used as the primary value for the methodological comparisons in this paper.

In the active-contact experiment, the median POI log_10_(*LR*) was 1.50, with values ranging from -1.32 to 3.10. Seventeen of the 20 active-contact cases gave POI likelihood ratios greater than 1. Thus, most active-contact cases supported *H*_1_, although the cases below 1 show that active contact does not guarantee prosecution-supporting evidence for the POI. The cases below 1 occurred where the POI quantity was low, or where an observed unknown contributor provided a plausible competing explanation for the direct-transfer role.

In the social-contact experiment, the POI likelihood ratio was never greater than 1. The median POI log_10_(*LR*) was -3.16, with values ranging from -6.24 to -1.30. Thus, all 20 social-contact cases supported *H*_2_ rather than *H*_1_ for the POI.

Observed unknown contributors sometimes received likelihood ratios greater than 1 in both experimental conditions. This is not contradictory. HaloGen evaluates each observed contributor as a possible direct actor, regardless of whether that contributor is named or unknown. A high LR for an unknown contributor indicates that the unknown contributor’s quantities are more compatible with the direct-transfer role than with secondary transfer. It does not imply support for the POI as the direct actor.

This behaviour is central to the method comparison. If an observed unknown contributor is treated explicitly, the contributor may compete with the POI for the direct-transfer role. If the same contributor is instead collapsed into a background term, the resulting POI LR may change. Selected illustrative cases are provided in Supplementary Table S1, and the full contributor-specific active-contact and social-contact results are provided in Supplementary Tables S2 and S3.

### 5.2 Comparison with simplified comparator approaches using Tippett plots

The behaviour of the likelihood ratios for the different approaches was evaluated using Tippett plots. Tippett plots, named after Tippett et al. [27], are commonly used as empirical performance diagnostics for LR systems [12, 21, 15], but they should be interpreted as dataset-specific checks of discrimination and misleading-evidence behaviour rather than as proof of universal calibration. A Tippett plot shows the empirical cumulative distribution of likelihood-ratio values from test cases with known ground truth. In this study, the x-axis is log_10_(*LR*), and the y-axis is the cumulative proportion of cases with LR values less than or equal to that value. The vertical line at log_10_(*LR*) = 0 corresponds to *LR* = 1, or neutral evidence. Values to the right of this line favour *H*_1_, while values to the left favour *H*_2_.

Figure 1 compares the HaloGen results with the discrete *M_x_* and binary source-support approaches. The upper panel shows the HaloGen Tippett curves for active-contact and social-contact ground-truth cases. The lower panel shows the discrete *M_x_*component-size approach for both active-contact and social-contact cases, together with the binary source-support approach for active-contact cases only.

**Figure 1:**
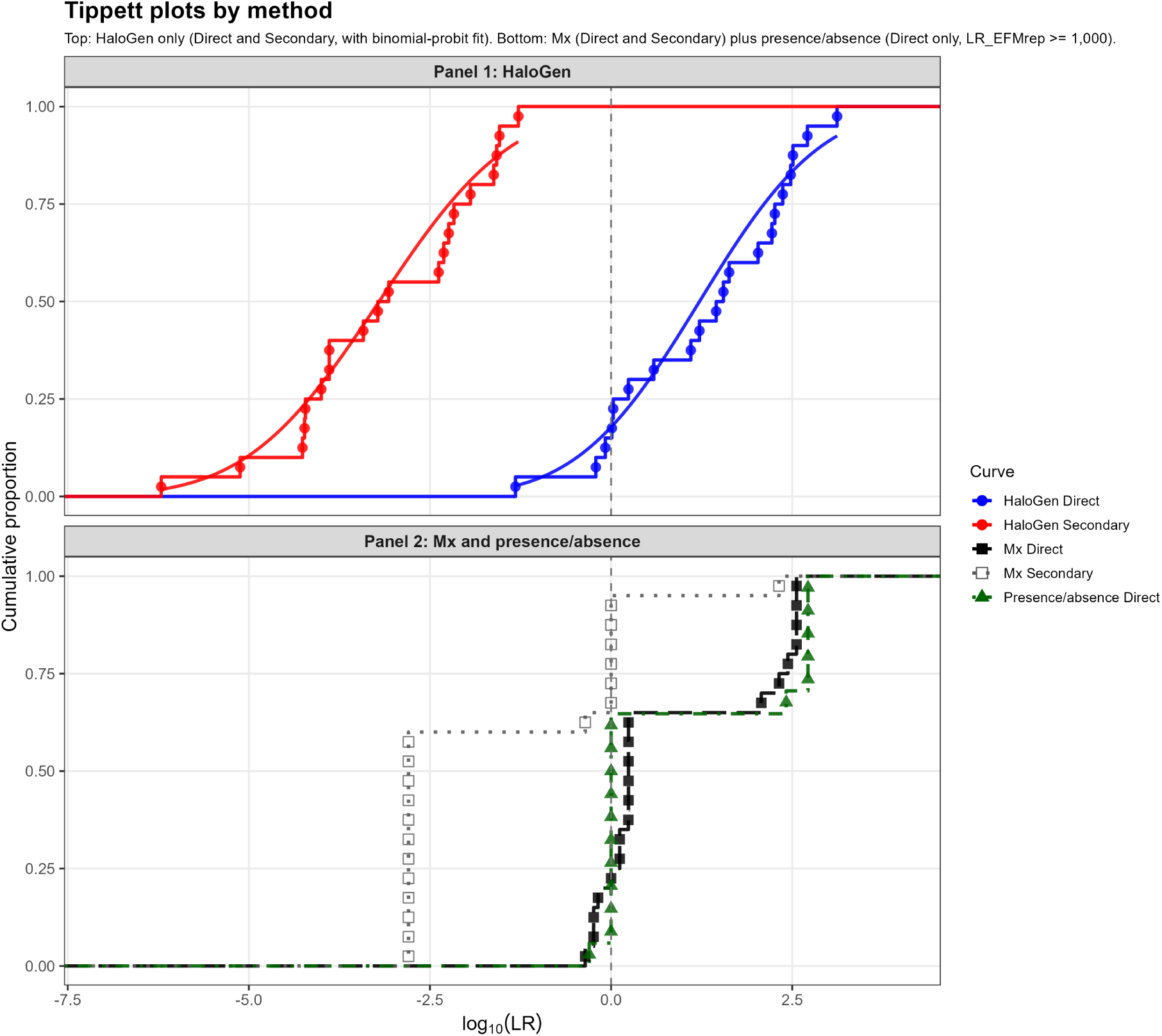
Comparison of activity-level LR distributions obtained using HaloGen and simplified comparator approaches. The upper panel shows HaloGen log_10_(*LR*) values for active-contact and social-contact ground-truth cases. The lower panel shows the discrete *M_x_* component-size approach for active-contact and social-contact cases, together with the binary source-support approach for active-contact cases. The discrete *M_x_* approach used *LR*_sub_*_−_*_source_ *>* 1 as a minimum source-support gate. Binary source support was defined using *LR*_sub_*_−_*_source_ *≥* 1000. No social-contact binary source-support curve is shown because no social-contact POI observation met this sub-source threshold. The dashed vertical line marks log_10_(*LR*) = 0, corresponding to neutral evidence.

The discrete *M_x_*approach used the minimum source-support restriction *LR*_sub_ *>* 1. This restriction should not be interpreted as a reporting threshold. It was applied because, at very low sub-source support, discrete *M_x_* estimates are unstable and the resulting absent/minor/major labels can be dominated by stochastic mixture-proportion variation. By contrast, the binary source-support approach used the more stringent threshold *LR*_sub_ *≥* 1000, representing a reportable source-level observation.

A social-contact binary source-support curve was not plotted because there were no social-contact samples above this threshold. The discrete *M_x_* approach retained more categorical information than the binary source-support approach, but still produced only limited separation between direct and secondary cases. In particular, some secondary-transfer discrete *M_x_* cases produced neutral or prosecution-supporting LRs. These points should be interpreted cautiously because the discrete *M_x_* approach is a categorical approximation built from thresholded component-size classes, not a full quantitative transfer model.

By contrast, the HaloGen Tippett curves were generated independently of any sub-source LR threshold. All activity-level cases were retained, including cases with low sub-source support.

### 5.3 Direct-transfer comparison with ReAct3 background approach

We next examined the effect of replacing explicit unknown-contributor assignment by a background/direct-unknown term. Figure 2 compares direct-transfer true cases using three methods: the full HaloGen calculation, the discrete *M_x_* component-size approach, and the ReAct3 background approach.

**Figure 2:**
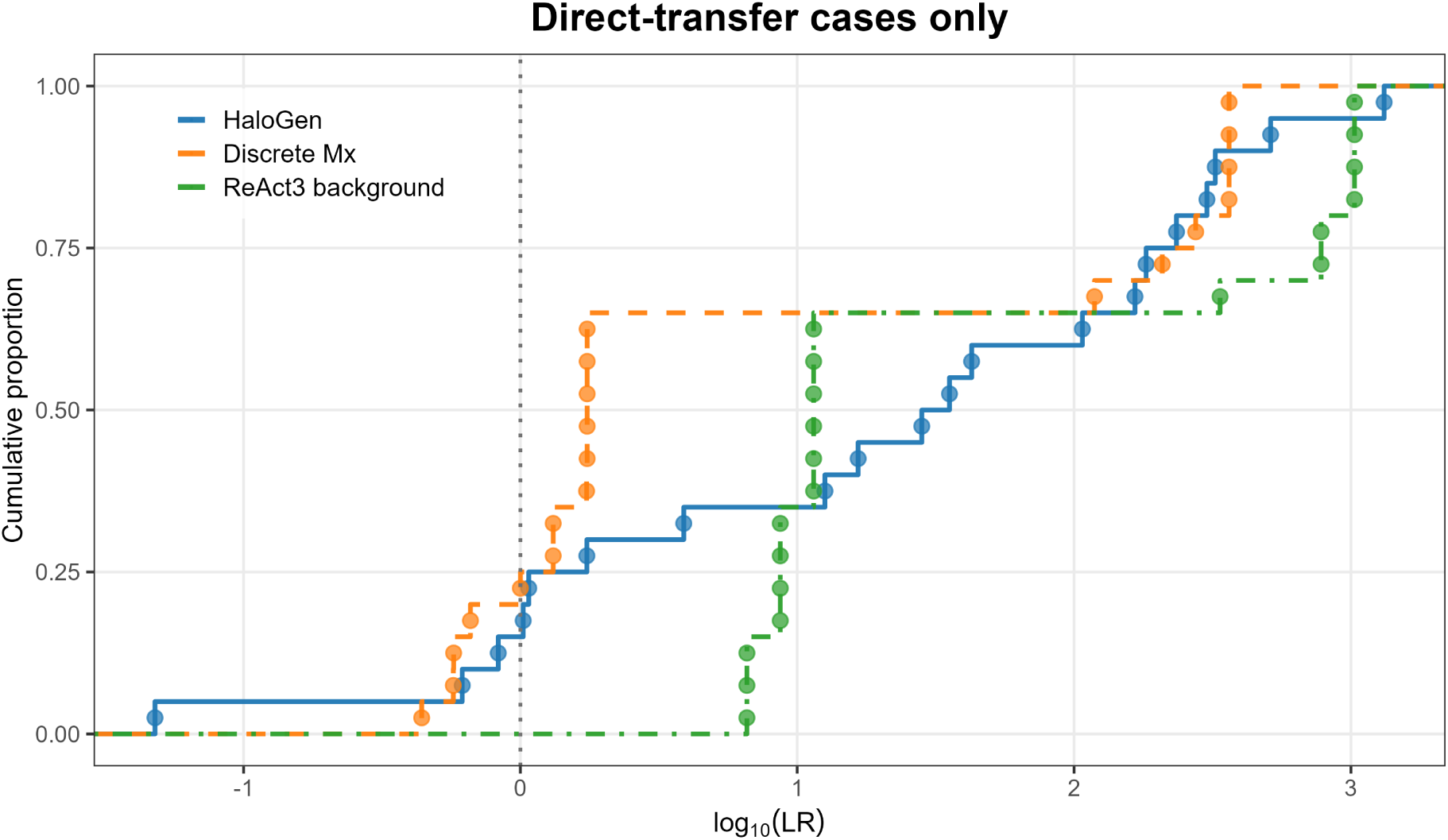
Direct-transfer cases only: comparison of HaloGen, discrete *M_x_*, and ReAct3 background approaches. The plot shows empirical cumulative distributions of log_10_(*LR*) for direct-transfer true cases. HaloGen uses the full contributor-specific quantitative information. The contributor-symmetric discrete *M_x_* approach allows observed unknown contributors to compete with the known POI for the direct-actor role. ReAct3 background uses the Experiment 3 background formulation. The ReAct3 background curve is shifted to the right of the discrete *M_x_*curve, indicating larger prosecution-supporting LRs for many direct-transfer cases.

The comparison was restricted to direct-transfer true cases because the social-contact samples had low sub-source LRs and would generally not be reported under a stringent source-level threshold. The direct-transfer cases therefore provide the most relevant comparison of how the approaches behave when the evidence is potentially reportable.

The ReAct3 background curve is shifted to the right of the discrete *M_x_* curve. This indicates that the background approach gives larger prosecution-supporting LRs for many direct-transfer cases. In the present data, the median shift of ReAct3 background relative to discrete *M_x_* approach was approximately 0.82 on the log_10_ scale, and 19 out of 20 direct-transfer cases had larger LRs under the ReAct3 background formulation than under the discrete *M_x_* formulation. Four direct-transfer cases crossed from defence-supporting or near-neutral values under the discrete *M_x_* calculation to prosecution-supporting values under the ReAct3 background calculation.

The comparison with HaloGen is also informative. The ReAct3 background curve was shifted to the right of HaloGen for part of the distribution. The median shift of ReAct3 background approach relative to HaloGen was approximately 0.44 on the log_10_ scale, and 12 out of 20 direct-transfer cases had larger LRs under ReAct3 background than under HaloGen.

These results show that the treatment of observed unknown contributors materially affects activity-level LRs. The ReAct3 background approach is useful as a simplified comparison, but it is not equivalent to a symmetry-preserving model. It answers a different modelling question because the observed unknown side is represented by a background/direct-unknown term rather than by evaluating each observed unknown contributor separately as a competing direct actor.

## 6 Discussion

### 6.1 Empirical discrimination between active and social contact

The HaloGen Tippett plot showed clear empirical discrimination between active-contact and social-contact cases in this dataset. Most active-contact cases gave LRs supporting *H*_1_, while all social-contact cases gave LRs supporting *H*_2_ for the POI. This supports the conclusion that contributor-specific DNA quantities, evaluated in an activity-level framework, can provide useful discrimination between active and social contact on stockings.

The use of the full-data Pop20-informed Lab-Bayes model for the Tippett plots also required consideration of data reuse. As described in Section 3.3, leave-one-case-out analysis was used to assess whether the local stockings observations contributing to calibration materially affected the LR calculated for the same case. The leave-one-case-out results were very close to the full-data results, and no case changed direction of support. The full-data Pop20-informed Lab-Bayes model was therefore retained as a convenient common reference model for comparing HaloGen with the simplified comparator approaches. The detailed leave-one-case-out procedure and results are provided in Supplementary Material S4.

The Tippett plot should not be overstated as proof of universal calibration. It is an empirical diagnostic based on known ground-truth cases from this particular dataset. It shows whether LRs tend to move in the expected direction and identifies misleading results, defined here as active-contact cases with *LR <* 1 or social-contact cases with *LR >* 1. External validation for other casework settings would require additional data generated under relevant conditions. Selected contributor-specific examples illustrating how the POI and observed unknown contributors were evaluated are provided in Supplementary Table S1, and the full active-contact and social-contact contributor-specific HaloGen results are provided in Supplementary Tables S2 and S3.

### 6.2 Information loss in discretised comparator approaches

The binary source-support and discrete *M_x_* approaches were useful benchmarks because they show what happens when continuous quantitative evidence is simplified to coarser summaries. The binary source-support approach was highly dependent on the chosen sub-source threshold. Using *LR*_sub_*_−_*_source_ *≥* 1000, the binary source-support approach produced a direct-transfer curve but no social-contact POI curve, because none of the social-contact POI observations met the threshold. The resulting social-contact binary source-support data would therefore consist only of threshold-negative observations and would not provide an informative Tippett curve.

The discrete *M_x_* approach retained more information because it distinguished absent, minor, and major component-size states. However, it remained a crude approach. It did not use the actual DNA quantities, and in some social-contact cases it produced neutral or prosecution-supporting LRs from observations with low sub-source support. This demonstrates the limitation of categorical approaches in low-level or complex mixture settings.

HaloGen performed better because it used the continuous contributor-specific quantities directly. It also retained all observations rather than requiring a source-level reporting threshold as a precondition for activity-level interpretation. The derivations of the binary source-support and discrete *M_x_* component-size approaches, including the Jeffreys-smoothed category probabilities and discrete contributor-assignment formulae, are provided in Supplementary Material S5.

### 6.3 Symmetry and observed unknown contributors

A central principle of activity-level interpretation is contributor-assignment symmetry. Suppose the evidence contains DNA from a named POI, *X*, and from an observed unknown contributor, *U* , and suppose there is one relevant direct actor. If the evidence does not distinguish *X* from *U* , the model should not favour *X* merely because *X* is named.

The two competing assignments are: *X* is the direct actor and *U* is a secondary contributor; or *U* is the direct actor and *X* is a secondary contributor. In a simple binary source-support example, let *t* be the probability of observing a contributor if that contributor is the direct actor, and let *s* be the probability of observing a contributor if that contributor is secondary. If both *X* and *U* are observed, the first assignment has likelihood *ts*, and the second has likelihood *st*. These are the same quantity, so the LR is 1. The evidence is neutral because it does not distinguish the named POI from the observed unknown contributor.

The same conclusion applies if the direct actor may also contribute through a secondary route, provided the same rule is applied to whichever contributor is assigned the direct role. The algebraic derivation is provided in Supplementary Material S5.

In HaloGen, the denominator is not restricted to a single generic unknown-background event. Instead, each observed unknown contributor is considered in turn as the possible direct actor under the defence proposition, while the POI is assigned to secondary transfer. HaloGen also includes the possibility that the true direct actor is not among the observed contributors, through the *F*_0_ non-detection term.

This symmetry acts as a safeguard against confirmation bias. If an observed unknown contributor is replaced by a single background parameter, the model no longer tests the unknown contributor as a possible direct actor. The unknown is effectively demoted from a competing explanation to a nuisance term. This can structurally favour the named POI, especially in mixtures where an observed unknown contributor is present at comparable or higher quantity. The Tengs case [9] provides an example where the judges ruled that confirmation bias had unfairly prevented the effect of unknown contributors being properly considered. HaloGen is designed to reduce this source of modelling bias. The same symmetry principle is demonstrated numerically in Supplementary Table S1, where selected cases show how observed unknown contributors can reduce the POI LR or receive a larger LR than the POI when their quantities are more compatible with the direct-transfer role. The corresponding simplified symmetry derivation is given in Supplementary Material S5, and the full HaloGen likelihood construction is given in Supplementary Material S2.

### 6.4 Why background approaches can shift LRs

Bayesian-network approaches [25, 3, 22] are valuable for structuring activity-level reasoning and for making assumptions explicit. However, the treatment of observed unknown contributors is a modelling decision that must be aligned with the propositions. In particular, where an observed unknown contributor may plausibly be the direct actor under the alternative proposition, replacing that contributor by a generic background term changes the proposition structure and is not equivalent to an exhaustive contributor-assignment model. The present paper demonstrates that the treatment of observed unknown contributors can materially affect the LR.

The ReAct3 background approach illustrates the practical effect of changing the treatment of observed unknown contributors. The ReAct3 formulation is not simply a model in which background cancels. It includes an unknown component under *H*_2_, represented by

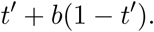

However, it still does not evaluate each observed unknown contributor separately as a possible direct actor.

The explicit formulae for the ReAct3 background approach used in this comparison are given in Supplementary Material S5.

In the direct-transfer cases, the ReAct3 background curve was shifted to the right of the discrete *M_x_* curve. This indicates that the background approach gave larger prosecution-supporting LRs for many report-relevant cases. The key reason is structural: the observed unknown contributor is no longer tested in the same contributor-specific way as the known relevant actor (POI). The background approach therefore changes the proposition structure.

This does not mean that background formulations are always inappropriate. They can be useful simplified models when the unknown component is genuinely background and not a plausible alternative actor. But it does imply that we need to think about how to assess background levels at the crime-scene. General experimental datasets may not fully capture item- or scene-specific background levels, because controlled transfer experiments are usually performed under restricted conditions. One possible approach is to obtain control samples from adjacent or comparable areas that are not directly associated with the alleged activity, so that item- or scene-specific background levels can be estimated rather than assumed. Such data would need to be incorporated transparently into the activity-level model and distinguished from observed contributors who may be plausible alternative actors. However, where an observed unknown contributor may be the direct actor under the alternative proposition, a symmetry-preserving assignment structure is preferable. The distinction between the ReAct3 background approach and the discrete *M_x_* component-size approach is therefore substantive. In the ReAct3 background approach, the observed unknown component is represented through a background/direct-unknown term. In the discrete *M_x_* component-size approach, observed unknown contributors are evaluated individually as possible direct actors under the alternative proposition. This difference in how the unknown contributor is modelled explains why the two approaches can give different LRs. The corresponding formulae are provided in Supplementary Material S5.

### 6.5 Role of Pop20-informed priors

The Pop20 compatibility diagnostics indicated that the stockings dataset was broadly compatible with the observed Pop20 reference range. The empirical upper-tail fractions were not small, and therefore did not indicate that the stockings dataset was an outlier. These diagnostics supported using Pop20 as regularising prior information, but not as a replacement for stockings-specific calibration data.

For this reason, the pure Pop20 Group model was not used as the primary result. Instead, Pop20 was used only through the Lab-Bayes model, where the Pop20-informed prior was updated by the local stockings observations. The detailed compatibility diagnostics, empirical upper-tail fractions, PCA plots and posterior predictive checks are provided in Supplementary Material S3.

The decisive sensitivity check was the weak-prior forced refit which did not rely upon any of the Pop20 datasets. This analysis used the same likelihood model, data structure, *F*_0_ policy, defined cases and case-level LR calculations as the primary analysis, but replaced the Pop20-informed prior with weakly informative hyperparameters that allowed substantially greater prior variance. The direct-transfer posterior was largely data-driven since it changed very little under the weak-prior refit. The secondary-transfer posterior showed greater prior sensitivity, as expected for the more variable and sparse secondary-transfer component, but this did not materially alter the case-level LR conclusions.

At the case level, the weak-prior medians fell within the corresponding Pop20-informed 10–90% posterior intervals for all defined cases. The median weak-prior minus Pop20-informed shift was small: approximately +0.21 log_10_ units for active-contact cases and +0.20 log_10_ units for social-contact cases. Where prior choice had a visible effect in active-contact cases, the Pop20-informed analysis tended to reduce prosecution-supporting LRs relative to the weak-prior analysis rather than inflate them.

Leave-one-case-out analysis gave the same practical conclusion. Omitting each activity-level case from the local calibration data and re-evaluating it using the fold-specific posterior produced only small shifts in log_10_(*LR*), and no case changed direction of support. Together, these findings support the use of the Pop20-informed Lab-Bayes model as a pragmatic regularised analysis for this single-laboratory stockings dataset.

### 6.6 Practical implications for casework

The results have several practical implications. First, observed unknown contributors should be considered explicitly whenever they cannot be eliminated as non-relevant actors. If an unknown contributor can be explained by a person with legitimate access to the victim or garment, such as a partner or household member, that person should be identified where possible and treated appropriately in the assessment. If the contributor remains unexplained, a symmetry-preserving activity-level method should allow that contributor to compete with the POI for the direct-transfer role.

Second, the sampling area should be defined before results are known. If multiple stains are collected and several contain no DNA of the POI, the LR for the POI may be reduced. Excluding such stains after observing the results risks confirmation bias. Predefining the relevant sampling area and inclusion criteria improves transparency and reduces the risk that interpretation is driven by selective use of favourable samples.

Third, simplified comparator approaches should be used cautiously. Binary source-support and discrete *M_x_* approaches are useful benchmarks, but they discard information. Background approaches may be appropriate in some settings, but they can alter the role of observed unknown contributors. The main advantage of HaloGen is that it evaluates the evidential contribution of each observed contributor explicitly, regardless of whether that contributor is named or unknown. For casework application, the worked formulae in Supplementary Material S2 and the simplified comparator derivations in Supplementary Material S5 provide useful checks on whether a proposed reporting model preserves contributor-level symmetry or instead changes the role assigned to observed unknown contributors.

## 7 Conclusion

This paper evaluated DNA transfer to stockings at activity level using HaloGen. The analysis showed that contributor-specific DNA quantities can discriminate between active-contact and social-contact ground-truth cases. HaloGen produced LRs in the expected direction for all social-contact cases and for most active-contact cases, while responding appropriately to low POI quantities and to observed unknown contributors that provided competing explanations for the direct-contact role.

Binary source-support and discrete *M_x_* approaches were useful as benchmarks but lost information relative to HaloGen. The ReAct3 background comparison showed that replacing explicit unknown-contributor assignment with a background/direct-unknown term can shift LRs upward in report-relevant direct-transfer cases. This supports the importance of preserving contributor-level symmetry in mixtures containing observed unknown contributors.

Because the stockings dataset was generated in a single laboratory, a Pop20-informed Lab-Bayes model was used. Compatibility and prior-sensitivity analyses supported this as a pragmatic regularisation strategy. The resulting framework provides a transparent method for evaluating activity-level propositions involving stockings, multiple stains, and observed unknown contributors.

## Supplementary material

**Supplementary Material S1: Data and contributor-specific HaloGen likelihood-ratio tables.** Contributor-specific quantity calculation, selected illustrative cases, and full Pop20-informed Lab-Bayes contributor-specific log_10_(*LR*) tables for active-contact and social-contact cases.

**Supplementary Material S2: HaloGen likelihood formulae.**Zero-augmented detectionlimit-aware lognormal transfer model, conditional-on-detection likelihoods, multiple-stain treatment, exhaustive elemental assignments, and *F*_0_ non-detection term.

**Supplementary Material S3: Pop20 compatibility diagnostics.** Robust Mahalanobis distance analysis, principal component analysis, posterior predictive transfer-rate checks, and diagnostic plots for the target stockings laboratory.

**Supplementary Material S4: Prior-sensitivity and leave-one-case-out diagnostics.** Weak-prior versus Pop20-informed prior comparison, posterior parameter summaries, case-level LR sensitivity, and leave-one-case-out recalibration results.

**Supplementary Material S5: Simplified comparator formulae.** Formula derivations for the binary source-support approach, discrete *M_x_* component-size approach, ReAct3 background approach, and contributor-symmetry checks.

**Supplementary Material S6: Reproducibility workflow.** Overview of the input workbooks, output files, cache objects, R programs and figure files used to reproduce the Tippett plots and method comparisons.

## Acknowledgements

Funding for this Project was received from the European Union’s Internal Security Fund -Police (ISFP) – with Grant Agreement title: Competency, Education, Research, Testing, Accreditation, and Innovation in Forensic Science” [CERTAIN-FORS] ISFP-2020-AG-IBA-ENFSI and number: 101051099. Also: ReAct II – Extension of ReAct. Work package 5 within the FOR-FUTURE project - Forensic Fundamentals, Technology, Multidisciplinarity, Research, Evaluation (FOR-FUTURE) — ISF-2023-TF2-AG-ENFSI-IBA-2.

