## Supplements for "Activity-level evaluation of DNA transfer to stockings: comparison of HaloGen with simplified contributor-based approaches"

**S6 Programs and resources** **21**

### S1 Data and contributor-specific HaloGen likelihood-ratio tables

This section provides the contributor-specific quantity calculations and likelihood-ratio tables supporting the main activity-level analysis.

#### S1.1 Contributor-specific quantities

Contributor-specific quantities were calculated as

$$Q_i = m_i Q_{\text{total}} E,$$

where $m_i$ is the mixture proportion assigned to contributor $i$, $Q_{\text{total}}$ is the total DNA concentration in the extract, and $E = 200\,\mu\text{L}$ is the elution volume. The activity-level dataset comprised 20 active-contact cases and 20 social-contact cases, each with two sampled stains. The female wearer was expected under both activity propositions and was not treated as a competing relevant actor.

#### S1.2 Selected HaloGen cases illustrating contributor-specific symmetry

Table S1 gives selected examples from the stockings dataset to illustrate contributor-specific HaloGen likelihood ratios. These examples are not intended to replace the full applied activity-level results reported in the companion experimental paper. They are included here to show how explicit assignment of observed unknown contributors can affect the POI likelihood ratio.

Table S1: Selected cases illustrating contributor-specific HaloGen likelihood ratios. Values are Pop20-informed Lab-Bayes median $\log_{10}(LR)$ values. Quantities are contributor-specific DNA quantities across the two sampled stains, in ng. In active-contact cases, the examples show how observed unknown contributors can reduce the POI LR when they provide plausible competing direct-actor assignments. In social-contact cases, the examples show that unknown contributors may receive LRs greater than 1 even when the POI LR supports $H_2$.

| Case | Context | POI quantities | POI $\log_{10}(LR)$ | Unknown quantities | Unknown $\log_{10}(LR)$ |
|---|---|---|---|---|---|
| F1_Dir | Active | 0.3278; 0.0878 | -1.32 | U1: 0.4172; 0.5221 | +1.30 |
| K1_Dir | Active | 1.6546; 0.7956 | -0.21 | U1: 3.5402; 1.8252 | +0.20 |
| J1_Dir | Active | 4.5920; 1.1088 | +0.23 | U1: 2.5830; 0.5040 | -0.24 |
| P1_Dir | Active | 2.3040; 1.0504 | +0.01 | U2: 1.1520; 1.5756 | -0.02 |
| K1_Sec | Social | 0; 0 | -5.12 | U1: 2.3684; 0.6816 | +3.42 |
| P1_Sec | Social | 0; 0 | -4.27 | U1: 0.3816; 0.3498 | +2.57 |
| I1_Sec | Social | 0.1847; 0.0810 | -2.16 | U1: 1.1286; 0.5670 | +2.10 |
| F1_Sec | Social | 0; 0 | -4.21 | U1: 0.1452; 4.3586 | +2.07 |
| M1_Sec | Social | 0; 0 | -3.89 | U1: 0.4488; 0.1704 | +1.87 |

The selected active-contact cases illustrate the symmetry-preserving behaviour of the model. In F1_Dir, the observed unknown contributor had larger quantities than the POI across the two stains and received the larger activity-level LR. In K1_Dir, the unknown contributor also had higher quantities than the POI and received an LR above 1, while the POI LR was below 1. In J1_Dir and P1_Dir, the POI quantities were substantial, but observed unknown contributors at comparable quantities reduced the POI LR toward neutrality.

The selected social-contact cases show the corresponding behaviour under $H_2$. The highest unknown-contributor LR was observed in K1_Sec, where the POI LR was strongly negative but unknown contributor U1 had a median $\log_{10}(LR)$ of $+3.42$. This distinction is important: the result does not support the POI as the direct actor. Rather, it indicates that another observed contributor may be more compatible with the direct-transfer role.

##### S1.3 Full contributor-specific HaloGen likelihood-ratio tables

Tables S2 and S3 give the full Pop20-informed Lab-Bayes median $\log_{10}(LR)$ results for the POI and observed unknown contributors. Missing unknown contributors are shown as "–".

Table S2: Full contributor-specific HaloGen results for active-contact cases.
Values are Pop20-informed Lab-Bayes median $\log_{10}(LR)$ values.

| Case | POI quantities | POI $\log_{10}(LR)$ | U1 quantities | U1 $\log_{10}(LR)$ | U2 quantities | U2 $\log_{10}(LR)$ |
|---|---|---|---|---|---|---|
| A1_Dir | 0.649; 2.3544 | +2.36 | 0.885; 0 | -2.59 | 0; 0.3924 | -2.93 |
| B1_Dir | 4.1488; 0.6372 | +2.71 | 0.4358; 0 | -2.85 | – | – |
| C1_Dir | 0; 0.1419 | -0.12 | – | – | – | – |
| D1_Dir | 5.3328; 2.9456 | +2.25 | 0.0606; 0.3682 | -2.38 | 0; 0.3682 | -3.19 |
| E1_Dir | 0.529; 0.8352 | +2.46 | 0.414; 0 | -2.61 | – | – |
| F1_Dir | 0.3278; 0.0878 | -1.32 | 0.4172; 0.5221 | +1.30 | – | – |
| G1_Dir | 0.196; 0.9048 | +0.03 | 0.224; 0.6264 | -0.04 | – | – |
| H1_Dir | 1.1856; 0.65 | +1.63 | 0.2736; 0.1 | -1.78 | 1.2768; 0 | -2.40 |
| I1_Dir | 0.9472; 1.4184 | +3.10 | 0; 0.197 | -3.42 | – | – |
| J1_Dir | 4.592; 1.1088 | +0.23 | 2.583; 0.504 | -0.24 | – | – |
| K1_Dir | 1.6546; 0.7956 | -0.21 | 3.5402; 1.8252 | +0.20 | 0; 0.1872 | -3.85 |
| L1_Dir | 0.8406; 0.1608 | +2.19 | 0.125; 0 | -2.70 | – | – |
| M1_Dir | 1.3216; 0.336 | +0.59 | 0.7552; 0.168 | -0.60 | – | – |
| N1_Dir | 1.449; 0.2496 | +2.49 | 0.1932; 0 | -2.81 | – | – |
| O1_Dir | 9.638; 5.472 | +1.21 | 0.366; 0.1847 | -1.22 | – | – |
| P1_Dir | 2.304; 1.0504 | +0.01 | 0.1472; 0.1212 | -2.58 | 1.152; 1.5756 | -0.02 |
| Q1_Dir | 2.1376; 1.2296 | +1.45 | 0.2004; 0.2968 | -1.46 | – | – |
| R1_Dir | 0.9956; 1.1011 | +2.02 | 0.283; 0.0823 | -2.05 | – | – |
| S1_Dir | 0.1881; 0.3053 | +1.09 | 0.1421; 0 | -2.28 | 1.0032; 0 | -1.22 |
| T1_Dir | 0.7569; 0.2436 | +1.54 | 0.0927; 0.1479 | -1.64 | 0; 0.1479 | -2.83 |

Table S3: Full contributor-specific HaloGen results for social-contact cases. Values are Pop20-informed Lab-Bayes median $\log_{10}(LR)$ values.

| Case | POI quantities | POI $\log_{10}(LR)$ | U1 quantities | U1 $\log_{10}(LR)$ | U2 quantities | U2 $\log_{10}(LR)$ |
|---|---|---|---|---|---|---|
| A1_Sec | 0.0672; 0.0234 | -1.95 | 0; 0.7527 | +0.73 | – | – |
| B1_Sec | 0; 0 | -2.25 | 0.2495; 0.0348 | +0.40 | – | – |
| C1_Sec | 0; 0 | -1.62 | 0.0265; 0.0294 | -1.81 | – | – |
| D1_Sec | 0.051; 0 | -1.30 | 0.0426; 0.0704 | -0.39 | – | – |
| E1_Sec | 0; 0 | -2.39 | 0.0704; 0.1184 | +0.59 | – | – |
| F1_Sec | 0; 0 | -4.21 | 0.1452; 4.3586 | +2.07 | 0; 0.2356 | -2.34 |
| G1_Sec | 0; 0 | -4.23 | 0.1554; 0.0635 | -1.83 | 0.2784; 0.4572 | +1.71 |
| H1_Sec | 0; 0.0016 | -6.24 | 0.0595; 0 | -0.90 | – | – |
| I1_Sec | 0.1847; 0.081 | -2.16 | 1.1286; 0.567 | +2.10 | – | – |
| J1_Sec | 0; 0 | -3.89 | 0.2516; 0.178 | -0.01 | 0.2516; 0.178 | -0.01 |
| K1_Sec | 0; 0 | -5.12 | 2.3684; 0.6816 | +3.42 | – | – |
| L1_Sec | 0.0094; 0 | -3.25 | 0.0643; 0 | -0.82 | – | – |
| M1_Sec | 0; 0 | -3.89 | 0.4488; 0.1704 | +1.87 | 0; 0.1704 | -2.24 |
| N1_Sec | 0.0094; 0 | -4.01 | 0.0547; 0.17 | +0.65 | – | – |
| O1_Sec | 0; 0 | -1.59 | – | – | – | – |
| P1_Sec | 0; 0 | -4.27 | 0.3816; 0.3498 | +2.57 | – | – |
| Q1_Sec | 0; 0.0881 | -2.31 | 0.1638; 0.2057 | +1.35 | 0.1638; 0 | -1.79 |
| R1_Sec | 0; 0 | -3.07 | 0.1649; 0.1282 | +1.36 | – | – |
| S1_Sec | 0; 0 | -3.42 | 0.3026; 0.1166 | +0.92 | 0; 0.5724 | -1.04 |
| T1_Sec | 0; 0.0296 | -1.66 | 0; 0.0099 | -3.11 | – | – |

#### S2 HaloGen likelihood formulae

HaloGen used zero-augmented, detection-limit-aware lognormal transfer models with parameters $(\mu, \sigma, k)$, where $k$ is the structural-zero or transfer-failure component. The detection limit was $DL = 0.001$ ng.

For detected quantities $q > DL$, the conditional-on-detection likelihood was

$$L(q \mid q > DL, \mu, \sigma) = \frac{f(q \mid \mu, \sigma)}{1 - F(DL \mid \mu, \sigma)},$$

where $f$ is the lognormal density and $F$ is the cumulative distribution function.

For an elemental assignment in which contributors in set $O$ are assigned to the direct role and the remaining observed contributors are assigned to secondary transfer, a simplified per-draw likelihood contribution has the form

$$\prod_{c \in O} t_c \prod_{c \notin O} s_c F_0^{N_S - |O|},$$

where $t_c$ and $s_c$ are contributor-specific direct and secondary likelihood terms, $N_S$ is the number of relevant direct actors, and $F_0$ accounts for required direct actors who are not detected among the observed contributors. In this study $N_S = 1$. Top-level proposition likelihoods are obtained by summing over admissible contributor-role assignments within each proposition.

#### S3 Pop20 compatibility diagnostics

This section gives the diagnostic calculations underlying the Pop20 compatibility statement in the main manuscript. The target data was the stockings; the reference set was the Pop20 cache set. These analyses were used as descriptive diagnostics of representativeness, not as formal hypothesis tests.

##### S3.1 Compatibility-assessment procedure

Having selected a Pop20-informed Lab-Bayes model, we assessed whether the target stockings laboratory was sufficiently related to the Pop20 reference data for regularisation to be reasonable. This assessment was intended as a representativeness and compatibility check, rather than as a test that the stockings laboratory was centrally representative of the reference dataset.

Specifically, we examined whether the target laboratory showed an anomalous direct-transfer distribution, secondary-transfer distribution, non-detection pattern, or direct–secondary ordering relative to the range of behaviour represented in the Pop20 reference data. The aim was to identify any structural discrepancy that would make Pop20-informed regularisation inappropriate.

Compatibility was assessed using posterior median parameter summaries extracted from stored HaloGen cache objects. For each laboratory, the direct-transfer channel was represented by three parameters,

$$(\mu_D, \log \sigma_D, \text{logit}(k_D)),$$

and the secondary-transfer channel by

$$(\mu_S, \log \sigma_S, \text{logit}(k_S)).$$

The structural-zero parameter $k$ was analysed on the logit scale because it is a probability constrained to the interval $(0, 1)$, whereas $\mu$ and $\log \sigma$ are on an unconstrained real scale.

Four feature representations were evaluated. The combined direct-plus-secondary representation was the six-dimensional vector

$$z_{\mathrm{D+S}} = (\mu_D, \log \sigma_D, \mathrm{logit}(k_D), \mu_S, \log \sigma_S, \mathrm{logit}(k_S)).$$

The direct-only and secondary-only representations were

$$z_D = (\mu_D, \log \sigma_D, \mathrm{logit}(k_D))$$

and

$$z_S = (\mu_S, \log \sigma_S, \mathrm{logit}(k_S)).$$

Finally, a contrast representation was used to describe the relationship between the secondary and direct transfer channels:

$$z_\Delta = (\Delta\mu, \Delta \log \sigma, \Delta\, \mathrm{logit}(k)),$$

where

$$\Delta\mu = \mu_S - \mu_D,$$

$$\Delta \log \sigma = \log \sigma_S - \log \sigma_D,$$

and

$$\Delta\, \mathrm{logit}(k) = \mathrm{logit}(k_S) - \mathrm{logit}(k_D).$$

The combined representation assesses whether the target laboratory is compatible with the reference laboratories in the absolute direct and secondary parameter space. The direct-only and secondary-only representations identify which channel contributes most to any apparent discrepancy. The contrast representation addresses a different question: whether the target laboratory preserves a direct–secondary relationship that is compatible with the reference data, even if its absolute parameter values lie toward the periphery of the reference set.

Robust Mahalanobis distances were computed for each feature representation using the minimum covariance determinant estimator. The target laboratory was compared with the Pop20 reference laboratories by calculating its robust Mahalanobis distance from the reference-data centre. These distances were used as descriptive multivariate diagnostics.

Principal component analysis and posterior predictive transfer-rate checks were used as complementary diagnostics. The posterior predictive check compared the target laboratory's posterior median probability of exceeding fixed quantity thresholds with the corresponding distribution from the 20-laboratory reference dataset for the direct and secondary channels.

#### S3.2 Compatibility results

The Pop20 compatibility diagnostics supported only a cautious use of Pop20-informed priors as regularising information for the stockings laboratory. Robust Mahalanobis distances were calculated for the combined direct-plus-secondary, direct-only, secondary-only and direct–secondary contrast representations. These were interpreted descriptively and together with the PCA and posterior predictive checks, rather than as formal outlier tests.

Table S4: Descriptive compatibility diagnostics for the target stockings laboratory relative to the Pop20 reference data. Robust Mahalanobis distances were computed using the minimum covariance determinant estimator. The empirical upper-tail fraction was calculated from the finite robust-distance reference distribution used for the diagnostic comparison. These values are descriptive compatibility diagnostics, not formal p-values.

| Feature representation | Dimension | $D^2$ | Empirical upper-tail fraction |
|---|---|---|---|
| Combined (D+S) | 6 | 217.1 | 0.2857 |
| Direct only | 3 | 114.0 | 0.2857 |
| Secondary only | 3 | 13.95 | 0.2857 |
| Contrast (S-D) | 3 | 5.431 | 0.4286 |

Table S4 shows that the stockings target data were not centrally representative of Pop20 in some absolute parameter representations. However, the empirical upper-tail fractions were not extreme, indicating that the target data were not outside the finite Pop20 compatibility range. The contrast representation gave the least separated result, with an empirical upper-tail fraction of 0.4286, supporting the interpretation that the direct–secondary relationship was not strongly discrepant from the Pop20 reference structure.

This distinction is important because the Pop20-informed Lab-Bayes analysis does not use the Pop20 data as a substitute for local stockings data. Instead, Pop20 provides regularising prior information, which is then updated by the local stockings observations. The compatibility diagnostics therefore argue against using a pure Pop20 Group model as the primary analysis, but they do not by themselves show that Pop20-informed regularisation materially affects the final case-level likelihood ratios. That question is addressed by the prior-sensitivity and leave-one-case-out analyses in Supplementary Material S4.

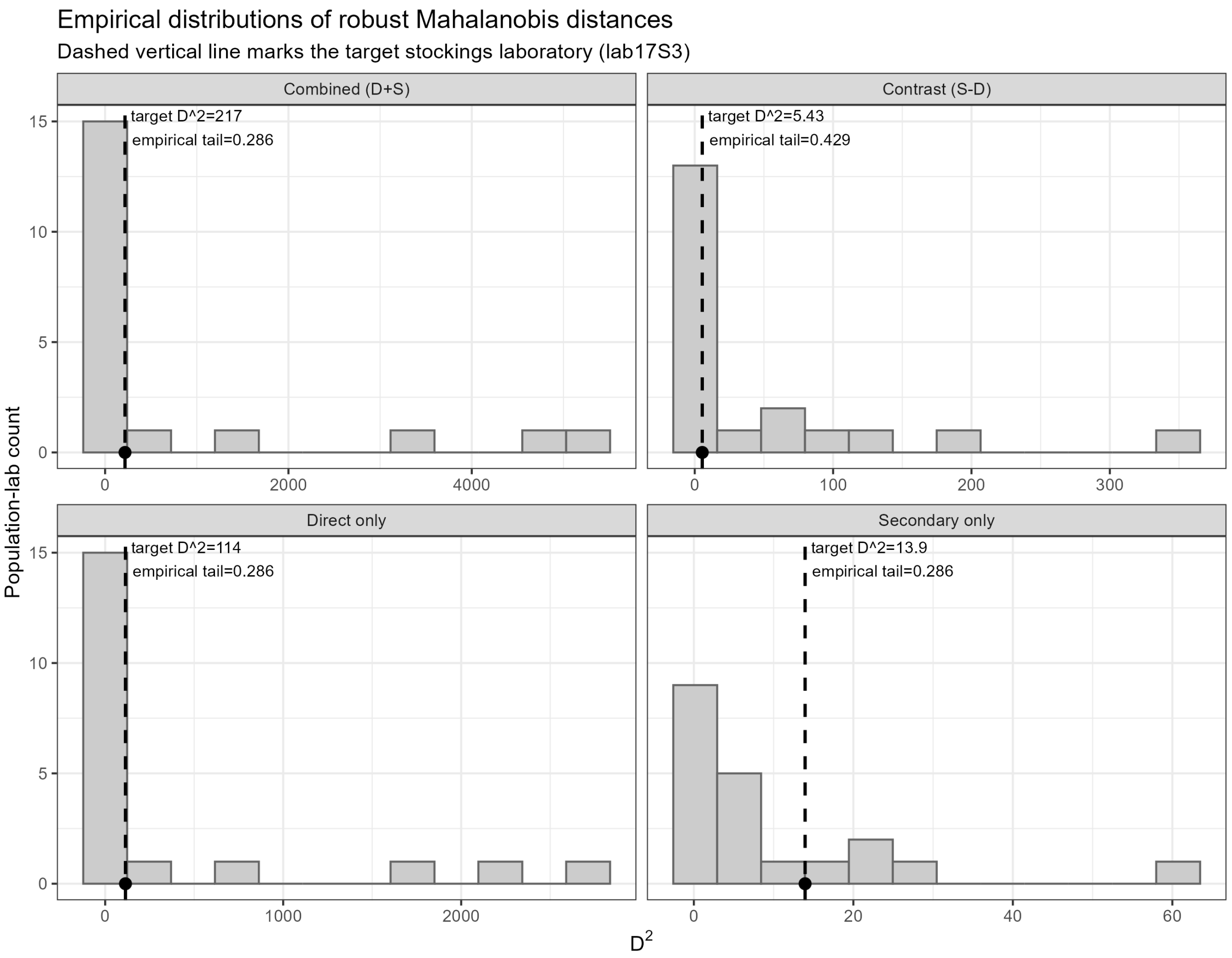

Figure S1: Empirical distributions of robust Mahalanobis distances for the Pop20 reference laboratories. The dashed vertical line marks `lab17S3`. These plots are descriptive finite-sample compatibility checks and should be interpreted together with the PCA plots and posterior predictive transfer-rate checks.

#### S3.3 PCA and posterior predictive checks

Principal component analysis was used as a visual diagnostic for the same four feature representations. In the combined representation, PC1 and PC2 explained 79.1% of the variance. In the direct-only, secondary-only and contrast representations, PC1 and PC2 explained 86.0%, 93.3% and 87.0% of the variance, respectively.

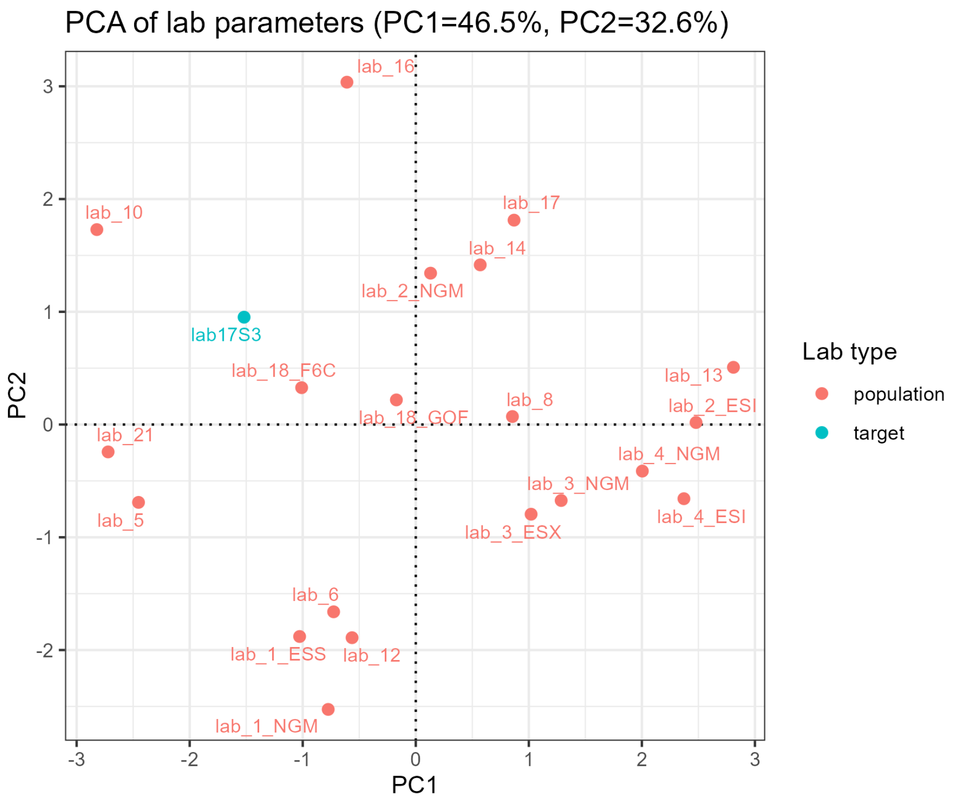

Figure S2: Principal component analysis of the combined direct-plus-secondary feature representation.

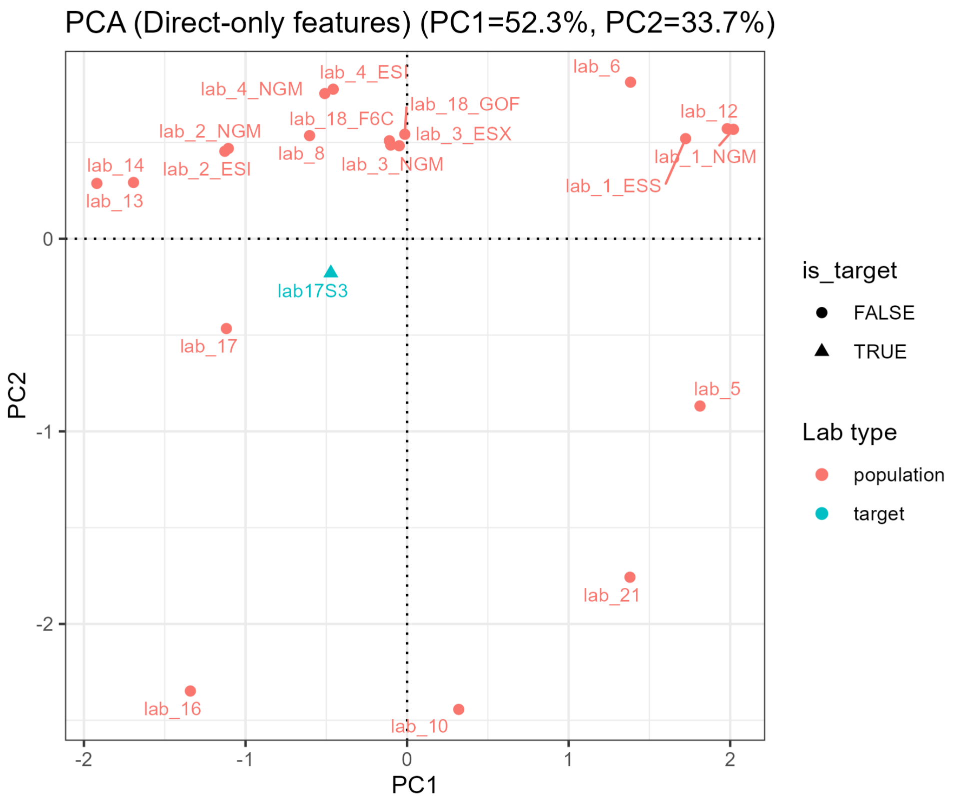

Figure S3: Principal component analysis of the direct-only feature representation.

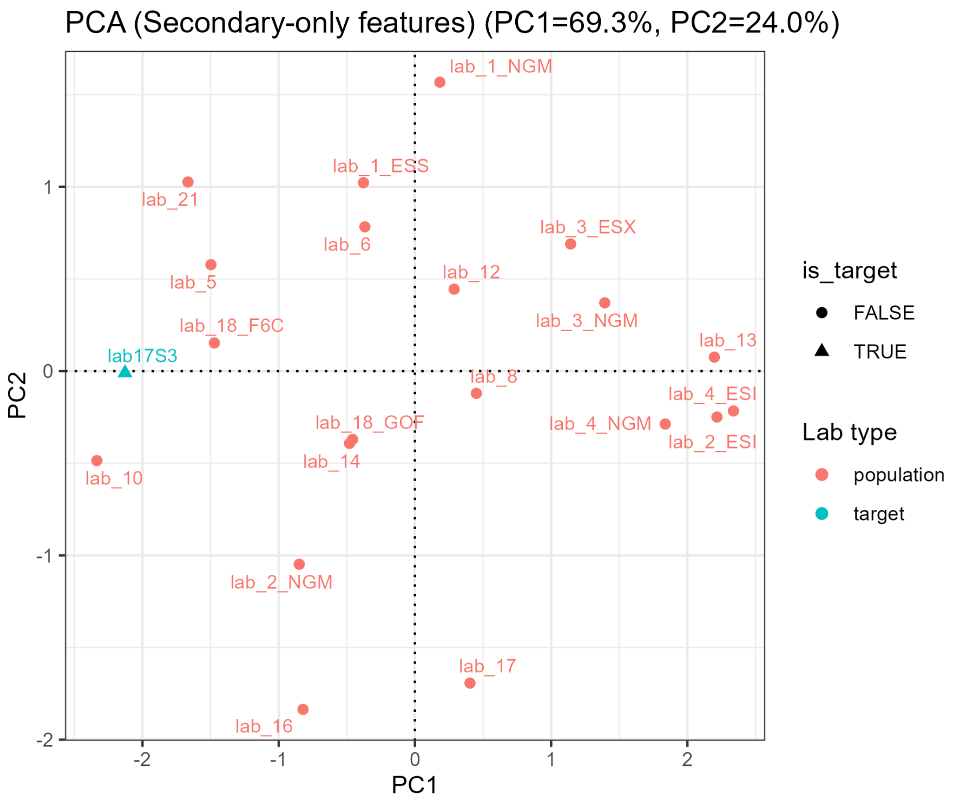

Figure S4: Principal component analysis of the secondary-only feature representation.

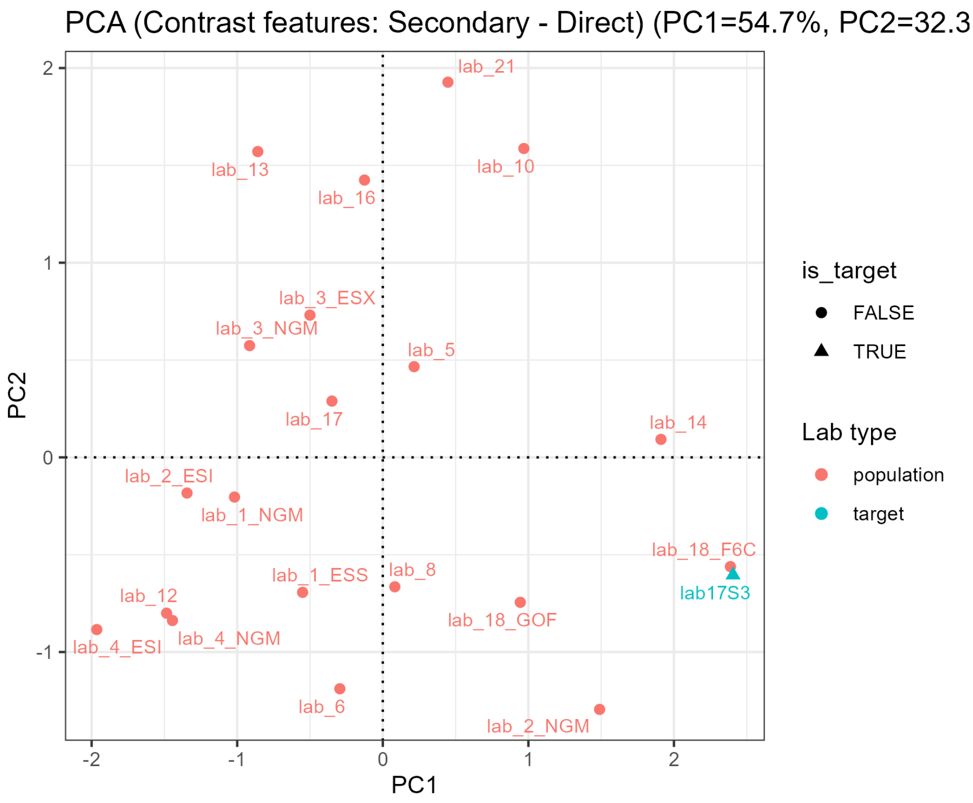

Figure S5: Principal component analysis of the contrast representation ($\Delta\mu, \Delta\log\sigma, \Delta\,\mathrm{logit}(k)$).

The PCA diagnostics were consistent with the robust-distance diagnostics. The target laboratory was peripheral in some absolute parameter representations, but the contrast representation showed better agreement with the Pop20 reference structure. This supports the interpretation that `lab17S3` should not be treated as centrally representative of Pop20, while still allowing cautious use of Pop20 as regularising prior information that is updated by local stockings data.

Posterior predictive transfer-rate checks compared the target laboratory's posterior median probability of exceeding fixed quantity thresholds with the Pop20 distribution. The direct channel was within the Pop20 10–90% band at all examined thresholds. The secondary channel was below the Pop20 10–90% band at several low thresholds and within the Pop20 range at higher thresholds. This channel-specific pattern is consistent with the decision to use Pop20 as regularising prior information rather than as a standalone Group model.

### S4 Prior-sensitivity and leave-one-case-out diagnostics

#### S4.1 Prior-sensitivity procedure and results

To evaluate the influence of prior specification, we compared the primary Pop20-informed Lab-Bayes analysis with a general weak-prior forced refit of the same Lab-Bayes model. This was not the production Lab-Vague model. The likelihood model, data structure, $F_0$ policy, defined cases and case-level likelihood-ratio calculations were kept unchanged; only the prior specification was varied.

The weak-prior model used weakly informative, neutral hyperparameters chosen to allow broad prior variance for both direct and secondary transfer:

$$\mu_0 = 0, \quad \tau_\mu = 10, \quad \log\sigma_0 = 0, \quad \tau_{\log\sigma} = 4, \quad \mu_k = 0.5, \quad \phi_k = 2.$$

The Pop20-informed model used channel-specific hyperparameters estimated from the Pop20 reference data and was used as the primary Lab-Bayes model in the main analysis.

Prior sensitivity was assessed at two levels. First, posterior summaries for the direct- and secondary-transfer parameters were compared under the weak-prior and Pop20-informed prior regimes. Second, the resulting case-level $\log_{10}(LR)$ values were compared for each defined case. In the case-level comparison, the Pop20-informed Lab-Bayes median and 10–90% posterior interval were treated as the reference, and the weak-prior Lab-Bayes median was compared against this interval.

Lab-Vague medians were also plotted as an additional sensitivity comparator, but the main prior-sensitivity comparison was between the Pop20-informed Lab-Bayes model and the weak-prior forced refit of the same Lab-Bayes model.

Table S5: Prior hyperparameters under weak and Pop20-informed prior regimes.

| | Direct transfer | | Secondary transfer | |
|---|---|---|---|---|
| Parameter | Weak | Pop20 | Weak | Pop20 |
| $\mu_0$ | 0 | -0.445 | 0 | -5.735 |
| $\tau_\mu$ | 10 | 1.830 | 10 | 2.968 |
| $\log\sigma_0$ | 0 | 0.452 | 0 | 1.017 |
| $\tau_{\log\sigma}$ | 4 | 0.585 | 4 | 0.525 |
| $\mu_k$ | 0.5 | 0.101 | 0.5 | 0.370 |
| $\phi_k$ | 2 | 7.131 | 2 | 3.654 |

Table S6: Posterior medians for laboratory-specific transfer parameters under weak and Pop20-informed prior regimes. The parameter $\sigma$ is obtained as $\exp(\log\sigma)$.

| | Direct transfer | | Secondary transfer | |
|---|---|---|---|---|
| Parameter | Weak | Pop20 | Weak | Pop20 |
| $\mu$ | -0.113 | -0.118 | -3.812 | -4.866 |
| $\log\sigma$ | 0.105 | 0.151 | 0.564 | 0.991 |
| $\sigma$ | 1.111 | 1.163 | 1.758 | 2.693 |
| $k$ | 0.041 | 0.030 | 0.711 | 0.630 |

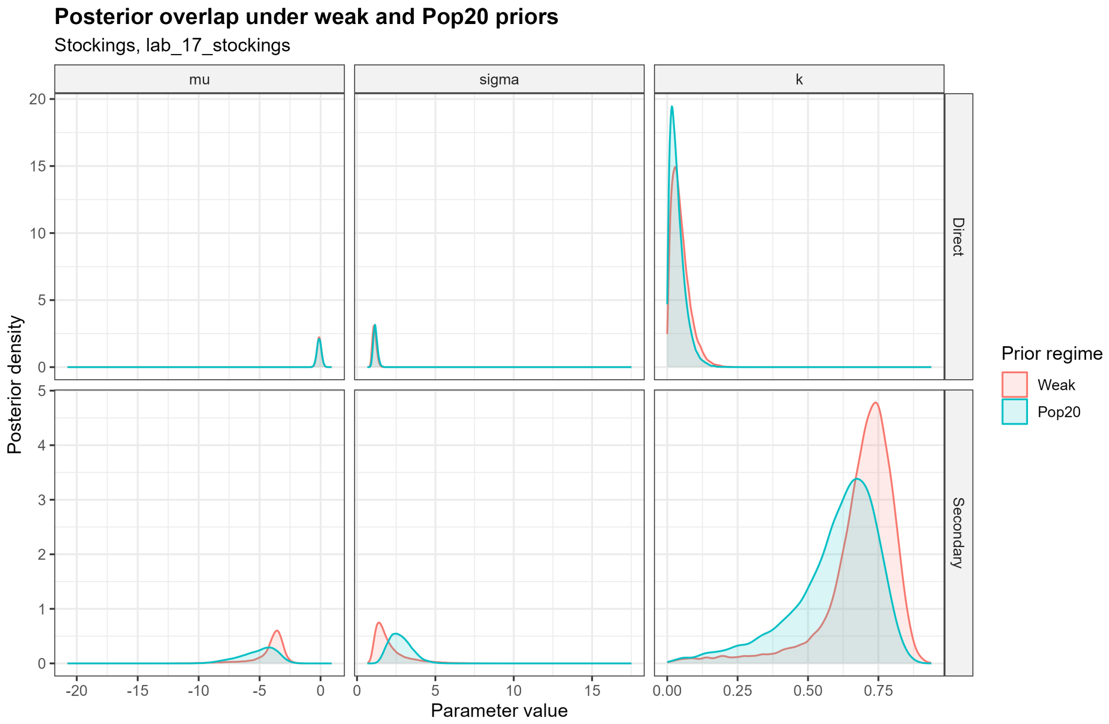

Figure S6: Posterior distribution overlap under weak and Pop20-informed prior regimes. Density curves show posterior distributions for $\mu$, $\sigma = \exp(\log \sigma)$, and $k$.

Table S7: Case-level sensitivity of Lab-Bayes $\log_{10}(LR)$ values to prior specification. Shift is weak-prior median minus Pop20-informed median.

| Arm | $n$ | Median shift | Min shift | Max shift | Inside Pop20 interval | Total |
|---|---|---|---|---|---|---|
| Active-contact | 20 | +0.21 | -0.50 | +1.52 | 20 | 20 |
| Social-contact | 20 | +0.20 | -0.90 | +0.58 | 20 | 20 |

Largest median shifts were O1_Dir (+1.52), D1_Dir (+1.12), A1_Sec (-0.90) and I1_Sec (-0.62). In all 40 cases, the weak-prior median fell within the corresponding Pop20-informed 10–90% posterior interval.

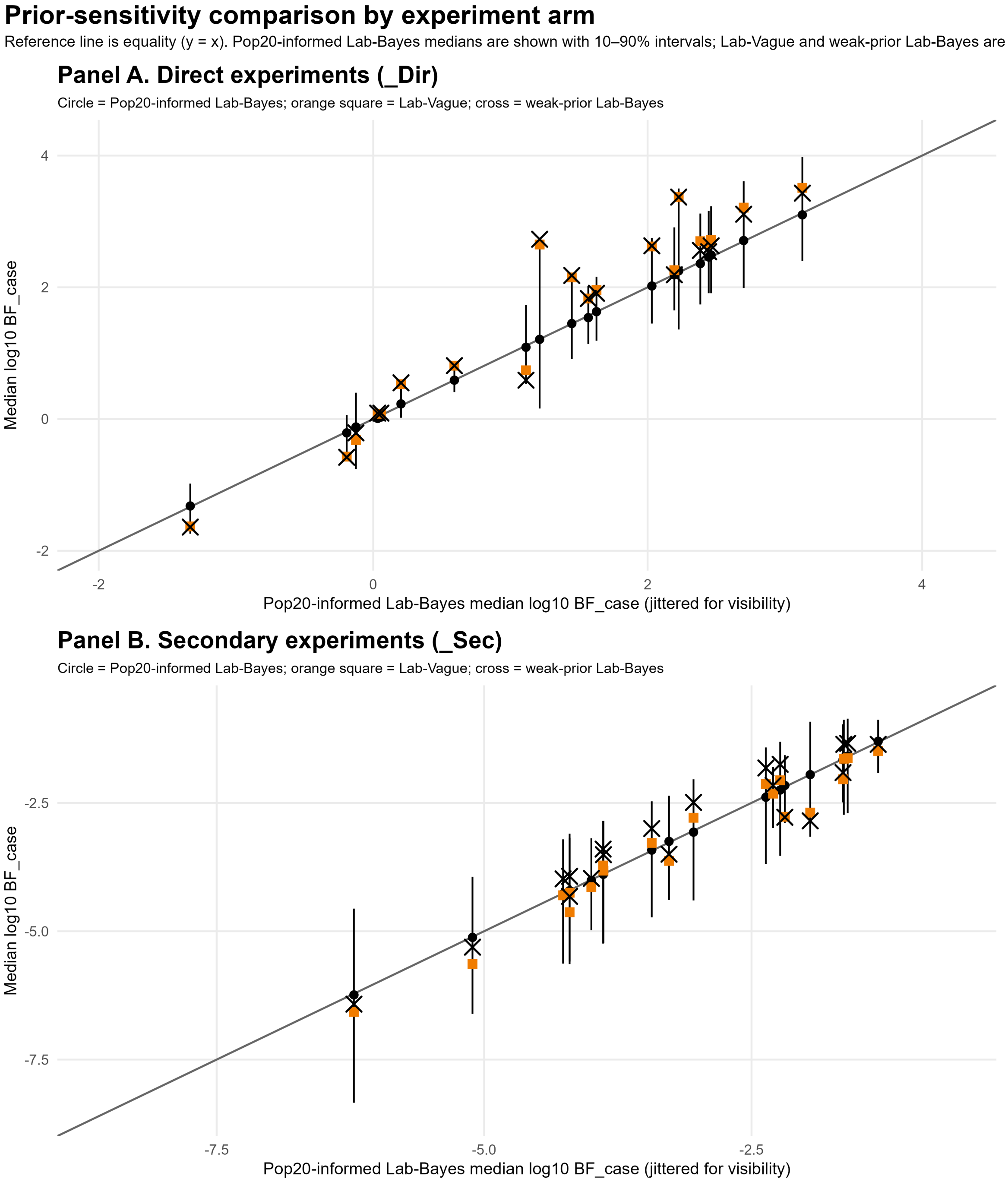

Figure S7: Case-level prior sensitivity for active-contact cases (Panel A) and social-contact cases (Panel B). The x-axis shows the Pop20-informed Lab-Bayes median $\log_{10}(LR)$. Black circles show the corresponding Pop20-informed Lab-Bayes medians on the y-axis, with vertical 10–90% posterior intervals. Orange squares show Lab-Vague medians, and black crosses show weak-prior Lab-Bayes forced-refit medians. The diagonal reference line represents equality with the Pop20-informed Lab-Bayes median. Small horizontal jitter is used only for visibility.

##### S4.2 Leave-one-case-out procedure and results

The leave-one-case-out analysis was performed to assess the effect of data reuse in the Tippett-plot and method-comparison analyses. The primary Tippett plots use the stockings cases as ground-truth test cases for assessing discrimination between active-contact and social-contact propositions. However, the same local stockings observations are also used to update the laboratory-specific Lab-Bayes transfer distributions. Thus, in the full-data analysis, a case can contribute both to the local calibration data and to the LR calculated for that same case.

The leave-one-case-out procedure circumvents this issue by omitting the case being evaluated from the local calibration data before calculating its LR. The omitted unit was the activity-level case, not an individual stain. Therefore, in each fold, both stain observations belonging to the held-out case were removed before refitting the Lab-Bayes model.

For active-contact cases, the two corresponding direct-transfer observations for the relevant actor/POI, denoted `LastH`, were removed from the local direct-transfer calibration data. For social-contact cases, the two corresponding secondary-transfer observations for the innocent/social-contact route, denoted `FirstH`, were removed from the local secondary-transfer calibration data. The Pop20-derived prior information was kept fixed in each fold because it was estimated from the external Pop20 reference data and was not part of the stockings experiment. The stockings-specific direct non-detection count used for the $F_0$ policy was updated within each fold.

This procedure provides a closer analogue of evaluating a case that did not contribute to the local calibration data. However, it is computationally intensive because a separate Lab-Bayes calibration fit is required for each held-out case. It also means that each case is evaluated using a slightly different local calibration dataset. For the main Tippett plots and simplified-comparator analyses, the full-data Pop20-informed Lab-Bayes calibration was therefore used as a convenient common reference model. The purpose of the leave-one-case-out analysis was to test whether this convenient full-data analysis materially affected the LR conclusions.

The resulting leave-one-case-out LRs were compared with the full-data Pop20-informed Lab-Bayes LRs.

Table S8: Leave-one-case-out sensitivity analysis for HaloGen Lab-Bayes activity-level likelihood ratios. Shift is LOO minus full-data Pop20-informed $\log_{10}(LR)$.

| Arm | $n$ | OK | Full-data | LOO | Shift | Min | Max | Crossings |
|---|---|---|---|---|---|---|---|---|
| Active-contact | 20 | 20 | 1.52 | 1.49 | -0.04 | -0.40 | +0.01 | 0 |
| Social-contact | 20 | 20 | -2.98 | -2.96 | +0.03 | -0.09 | +0.15 | 0 |

The leave-one-case-out results were very close to the full-data Pop20-informed results. For active-contact cases, the median $\log_{10}(LR)$ changed from 1.52 to 1.49. For social-contact cases, the median changed from $-2.98$ to $-2.96$. No case changed direction of support after leave-one-case-out recalibration. These results indicate that the separation shown in the Tippett plots was not materially caused by the case under evaluation also contributing to the local calibration data.

#### S5 Simplified comparator formulae

##### S5.1 Derivation of simplified comparator likelihood ratios

This section gives the derivations for the simplified comparator approaches used in the main text. These approaches were not intended to reproduce the full HaloGen calculation. They were used to examine the consequences of reducing contributor-specific quantitative DNA information to simpler binary or categorical summaries.

Throughout this section, the activity-level propositions are:

$$H_1 : \text{the POI is the direct actor,}$$

and

$$H_2 : \text{the POI is not the direct actor.}$$

The number of relevant direct actors is $N_S = 1$. Observed unknown contributors are not automatically treated as background. Under $H_2$, an observed unknown contributor may be the direct actor, or the relevant direct actor may be unobserved.

For these simplified calculations, $x_i = (x_{i1}, \ldots, x_{iR})$ denotes the simplified observation vector for contributor $i$ across the $R$ sampled stains. In the binary source-support approach, $x_{ir} \in \{0, 1\}$. In the discrete $M_x$ component-size approach, $x_{ir} = C_{ir} \in \{\text{Absent}, \text{Minor}, \text{Major}\}$. Thus $x_i$ is generic notation for the simplified observation; it is not a continuous quantity.

Let $\mathcal{U} = \{U_1, \ldots, U_m\}$ denote the set of observed unknown contributors. Let $D_i = \Pr(x_i \mid \text{direct transfer})$ and $S_i = \Pr(x_i \mid \text{secondary transfer})$. Also define

$$D_0 = \Pr(\text{direct actor unobserved on all } R \text{ stains}),$$

the simplified analogue of the HaloGen $F_0$ non-detection term.

Under $H_1$, the POI is assigned to the direct role and all observed unknown contributors are assigned to the secondary role:

$$L(H_1) = D_{\text{POI}} \prod_{u \in \mathcal{U}} S_u.$$

Under $H_2$, the POI is assigned to the secondary role. The relevant direct actor may be one of the observed unknown contributors, or may be unobserved:

$$L(H_2) = S_{\text{POI}} \left[ D_0 \prod_{u \in \mathcal{U}} S_u + \sum_{u \in \mathcal{U}} D_u \prod_{\substack{v \in \mathcal{U} \\ v \neq u}} S_v \right].$$

The resulting POI likelihood ratio is

$$LR_{\text{POI}} = \frac{D_{\text{POI}} \prod_{u \in \mathcal{U}} S_u}{S_{\text{POI}} \left[ D_0 \prod_{u \in \mathcal{U}} S_u + \sum_{u \in \mathcal{U}} D_u \prod_{\substack{v \in \mathcal{U} \\ v \neq u}} S_v \right]}.$$

This is the simplified contributor-symmetric LR formula for a binary or categorical representation of the observed evidence.

#### S5.2 Binary source-support approach

For the binary source-support approach, the generic simplified observation $x_{ir}$ was defined as a binary indicator:

$$x_{ir} \in \{0, 1\},$$

where $x_{ir} = 1$ denotes a supported contributor assignment for contributor $i$ on stain $r$, and $x_{ir} = 0$ denotes no supported contributor assignment.

This was deliberately not a quantity threshold based only on $Q_i > DL$. Presence was defined using a sub-source support threshold,

$$LR_{\text{sub}} \geq 1000,$$

where $LR_{\text{sub}}$ denotes the sub-source likelihood ratio obtained from EFMrep. This threshold was intended to approximate a reportable source-level observation.

Let $p_D = \Pr(x = 1 \mid \text{direct transfer})$ and $p_S = \Pr(x = 1 \mid \text{secondary transfer})$. The direct-transfer presence count was 35 out of 40 and the secondary-transfer presence count was 0 out of 40. Jeffreys correction was applied, giving

$$\widehat{p}_D = \frac{35 + 1/2}{40 + 1} = 0.8659,$$

and

$$\widehat{p}_S = \frac{0 + 1/2}{40 + 1} = 0.0122.$$

For contributor $i$, the direct and secondary binary source-support likelihoods across $R$ stains were

$$D_i = \prod_{r=1}^{R} \widehat{p}_D^{x_{ir}} (1 - \widehat{p}_D)^{1 - x_{ir}},$$

and

$$S_i = \prod_{r=1}^{R} \widehat{p}_S^{x_{ir}} (1 - \widehat{p}_S)^{1 - x_{ir}}.$$

The probability that an unobserved direct actor is absent across all $R$ stains was

$$D_0 = (1 - \widehat{p}_D)^R.$$

Substitution of these terms into the contributor-symmetric assignment expression above gives the binary source-support POI likelihood ratio.

#### S5.3 Discrete $M_x$ component-size approach

For the discrete $M_x$ approach, each contributor on each stain was classified into one of three categories:

$$C_{ir} \in \{\text{Absent}, \text{Minor}, \text{Major}\}.$$

Major/minor terminology is commonly used in forensic DNA mixture interpretation, but there is no universal mixture-proportion threshold defining a major contributor. A threshold of approximately 0.6 has previously been used as an operational definition of a major contributor in a related transfer and activity-level study [1]. The same threshold was used here only as an operational category for the simplified comparator. It was chosen to represent a clear majority component in this controlled comparison. It is deliberately restrictive and should not be interpreted as a general standard. The threshold is easiest to interpret in simple two-contributor settings, but the present analysis includes observed unknown contributors; for this reason, the threshold should be regarded only as an operational category for the simplified comparator.

A contributor was classified as absent if no DNA quantity was attributed to that contributor. A detected contributor was classified as major if $M_x \geq 0.6$, and as minor if $0 < M_x < 0.6$.

The discrete $M_x$ approach used the minimum sub-source support restriction

$$LR_{\text{sub}} > 1.$$

This was not a reporting threshold. It was used because, at very low sub-source support, estimated mixture proportions can be unstable and the resulting absent/minor/major categories can be dominated by stochastic mixture-proportion variation. Contributor-stain combinations with no attributed quantity were retained as absent observations for the purposes of estimating the absent category and the simplified non-detection term.

Let $\pi_D(c) = \Pr(C = c \mid \text{direct transfer})$ and $\pi_S(c) = \Pr(C = c \mid \text{secondary transfer})$, for $c \in \{\text{Absent}, \text{Minor}, \text{Major}\}$. The category probabilities were estimated from the retained observations separately for the direct and secondary transfer datasets. With Jeffreys smoothing for the three-category distribution,

$$\widehat{\pi}_D(c) = \frac{n_D(c) + 1/2}{n_D + 3/2},$$

and

$$\widehat{\pi}_S(c) = \frac{n_S(c) + 1/2}{n_S + 3/2},$$

where $n_D(c)$ and $n_S(c)$ are the retained direct and secondary counts in category $c$, and $n_D$ and $n_S$ are the corresponding retained totals.

For contributor $i$, the direct and secondary categorical likelihoods across $R$ stains were

$$D_i = \prod_{r=1}^{R} \widehat{\pi}_D(C_{ir}),$$

and

$$S_i = \prod_{r=1}^{R} \widehat{\pi}_S(C_{ir}).$$

The probability that an unobserved direct actor is absent across all $R$ stains was

$$D_0 = \{\widehat{\pi}_D(\text{Absent})\}^R .$$

Substitution of these terms into the contributor-symmetric assignment expression above gives the binary source-support POI likelihood ratio.

#### S5.4 Symmetry check

The simplified contributor-symmetric assignment structure preserves contributor symmetry. If one named POI and one observed unknown contributor have the same simplified observation state, then the direct and secondary likelihood terms are the same for the two contributors. The comparison between assigning the direct role to the POI and assigning it to the unknown contributor is therefore neutral:

$$\frac{D_{\text{POI}} S_U}{S_{\text{POI}} D_U} = 1 \quad \text{when} \quad D_{\text{POI}} = D_U,\ S_{\text{POI}} = S_U.$$

This is the same symmetry principle implemented more generally in HaloGen using continuous quantitative likelihoods.

#### S5.5 ReAct3 background approach

The ReAct3 background approach was used as a simplified comparator to examine the effect of replacing explicit unknown-contributor assignment by a background/direct-unknown term.

Let

$$t = \Pr(\text{POI observed} \mid \text{direct transfer}),$$

$$s = \Pr(\text{POI observed} \mid \text{secondary transfer}),$$

$$t' = \Pr(\text{unknown direct actor observed} \mid H_2),$$

and

$$b = \Pr(\text{background unknown DNA observed}).$$

For a single stain where only the POI is observed, the approach used was

$$LR_{\text{POI}} = \frac{t}{s(1-t')}.$$

Where both the POI and an unknown contributor are observed, the approach used was

$$LR_{\text{POI+U}} = \frac{tb}{s\{t' + b(1-t')\}}.$$

In the present analysis, the background parameter was set to $b = 0.3$.

This approach differs from the binary source-support and discrete $M_x$ calculations because the observed unknown contributor is not evaluated individually as a possible direct actor. Instead, the unknown side is represented by a background/direct-unknown term. The ReAct3 background approach therefore answers a different modelling question from the contributor-symmetric binary source-support and discrete $M_x$ calculations.

#### S6 Programs and resources

The current analysis used the Pop20-informed Lab-Bayes cache file `lab17S3.rds` stockings dataset, available at `https://github.com/peterdgill/HaloGen_Bayes/blob/main/stockings`. Halogen v.3.0.0 was used for the analysis: `https://github.com/peterdgill/HaloGen_Bayes/releases/tag/v3.0.0`
